# Mutations in the HKD motif of *Trypanosoma brucei* cardiolipin synthase inhibit cardiolipin synthesis and parasite growth

**DOI:** 10.64898/2025.12.15.694347

**Authors:** Advaitha Iyer, Peter Bütikofer

## Abstract

Phospholipases D (PLDs) are ubiquitous enzymes of the PLD superfamily that catalyze phosphodiester bond cleavage and transphosphatidylation reactions. They are characterized by the presence of two conserved HXK(X)_4_D(X)_6_G(X)_2_N or HKD motifs. While these motifs are essential for catalysis in diverse PLD family members, their functional significance in the bacterial-type cardiolipin synthase of the protozoan parasite Trypanosoma brucei (TbCls) has not yet been investigated.

TbCls is essential for parasite survival in culture and contains two conserved HKD motifs in its amino acid sequence. Here we take advantage of a previously constructed cell line in which the endogenous alleles of TbCls were replaced with an inducible ectopic copy of the enzyme, to introduce point mutations in the HKD motifs of TbCls. We found that the HKD motif and its surrounding amino acids are essential for the de-novo synthesis of cardiolipin and for growth of the parasite.

## Introduction

Phospholipases D (PLDs) are ubiquitous enzymes belonging to the PLD superfamily and have been studied in bacteria, fungi, mammals and plants(1). PLDs participate in hydrolysis and transphosphatidylation reactions, by cleaving phosphodiester bonds of glycerophospholipids to generate phosphatidic acid (PA) and a free hydrophilic head group, and transphosphatidylating phosphatidyl groups to acceptor primary alcohols, respectively(2). PLD enzymes are involved in diverse biological functions including lipid biosynthesis, receptor signaling pathways and cytoskeletal morphology(3),(4). In addition, they play important roles in diseases like diabetes by regulating insulin signaling(5), certain types of cancer by driving cell proliferation and migration(6–8), and Alzheimer’s disease and Parkinson’s disease by mediating amyloid neurotoxicity and accumulating in Lewy bodies, respectively(4, 9–11).

Members of the PLD superfamily are characterized by the presence of highly conserved HXK(X)_4_D(X)_6_G(X)_2_N (henceforth HKD) motifs in their primary amino acid sequences. HKD motifs are also present in other enzymes involved in glycerophospholipid turnover, such as bacterial-type cardiolipin synthases (Cls), phosphatidylserine synthases (PSS), bacterial endonucleases and certain viral proteins(12–15). Several mutagenesis studies have revealed the importance of the HKD motif for catalytic activity. Mutation of any conserved amino acid in either of the two HKD motifs of human PLD1 resulted in complete loss of enzymatic activity(13), supporting earlier predictions that the HKD motif is critical for catalysis(16–18) and that the two motifs do not function independently(13). In rat PLD, mutations in the HKD motif not only caused the loss of enzymatic activity but also disturbed the association of the N- and C-terminal domains(19). Furthermore, replacement of the histidine residue in either of the two HKD motifs significantly lowered *V. parahaemolyticus* PSS activity and abolished *E. coli* ClsC activity(12, 20). Finally, point mutations in the HKD motif of F13L protein of vaccinia virus resulted in diminished plaque formation and loss of viral cellular egress(13, 14).

The crystal structure of *A. thaliana* PLD1 revealed that the two HKD subdomains are contiguous and form the core of the catalytic domain(18). In addition, the two HKD domains were found to share structural and topological similarities, with each domain exhibiting a three-layered sandwich conformation containing 7 β-strands flanked by 3 or 1 α-helices, respectively, with the α and β structures being connected by loops. The N-terminal C2 domain showed exclusive interaction with the second HKD domain(18). Based on mutagenesis and structural studies, a two-step catalytic mechanism was proposed for enzymes of the PLD superfamily. The mechanism predicts the histidine residue of the first HKD subdomain to act as a nucleophile, leading to the formation of a phosphoenzyme intermediate followed by the release of the head group. The histidine in the second HKD subdomain would act as an acid to propagate the release of PA(16, 18). All PLDs containing the two HKD motifs are believed to follow a similar catalytic mechanism.

*Trypanosoma brucei* is a eukaryotic protozoan parasite that causes African sleeping sickness in humans and nagana in domestic animals. Trypanosomes *de novo* synthesize lipids and exhibit phospholipid compositions resembling other eukaryotes, with phosphatidylcholine and phosphatidylethanolamine being the most abundant phospholipid classes, followed by phosphatidylinositol, phosphatidylserine and cardiolipin (CL) as minor phospholipid classes(21–23). Interestingly, and in contrast to most other eukaryotes, the trypanosome genome encodes a bacterial-type cardiolipin synthase, having two conserved HKD motifs and belonging to the PLD superfamily(24). Bacterial-type cardiolipin synthases catalyze the last step in CL synthesis using phosphatidylglycerol (PG) and a phosphatidyl moiety of another PG as substrates. In contrast, eukaryotic-type cardiolipin synthases use PG and CDP-diacylglycerol as substrates for CL formation(25). The expression of *T. brucei* cardiolipin synthase (TbCls) is essential for survival of both *T. brucei* procyclic(26) and bloodstream(27) forms in culture. The role of the HKD motifs in TbCls enzyme function has not been studied.

In this study, *T. brucei* procyclic forms, in which the endogenous alleles of TbCls have been deleted and an inducible ectopic copy of the enzyme has been introduced to maintain parasite growth(26), are used to investigate the importance of the HKD motifs and selected surrounding amino acids for TbCls enzymatic activity and CL synthesis in vivo and parasite growth in culture. We found that point mutations in either of the two HKD motifs decreased CL production and resulted in growth inhibition of parasites. Mutations of the histidine residue in the first HKD motif caused a more severe growth defect than mutations of the lysine residue. Our results also point to the significance of small nonpolar amino acids adjacent to the HKD motif for TbCls activity, CL synthesis and parasite growth.

## Results and Discussion

### Point mutations in the HKD motif reduce parasite growth

To study the importance of the two conserved HKD motifs for TbCls function in vivo, conditional TbCls knockout parasites expressing a tetracycline-inducible HA-tagged copy of wild-type TbCls (TbCls-HA) were complemented with constitutively expressed cMyc-tagged wild-type TbCls (as proof of principle) or TbCls enzymes carrying point mutations in one or both HKD domains, or in a conserved region flanking the second HKD domain. The TbCls mutations are indicated in Fig. 1. Since *T. brucei* procyclic forms depend on TbCls function for CL synthesis and parasite survival in culture(26), reduced growth of trypanosomes after removal of tetracycline from the culture medium (to ablate TbCls-HA expression) was taken as indication for decreased, or absent, TbCls activity.

**Fig. 1.**
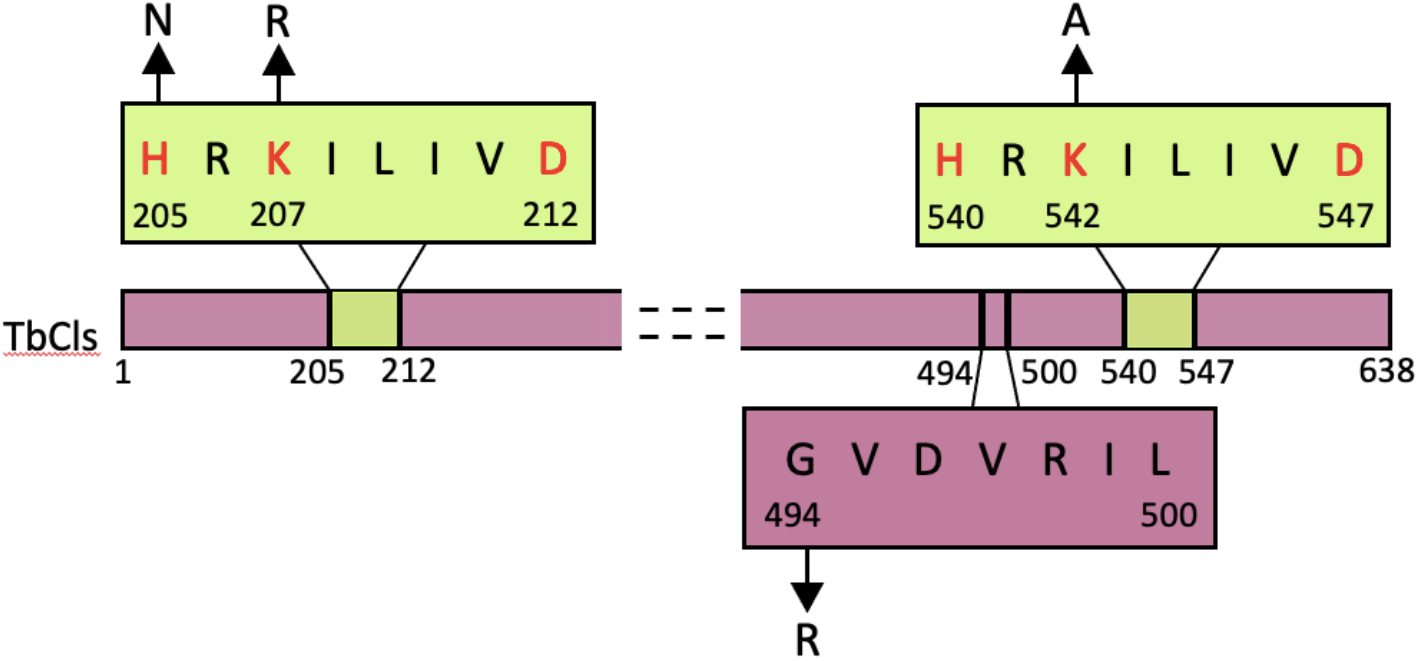
Amino acid sequences of the two HKD motifs in *T. brucei* cardiolipin synthase (TbCls). The minimal PLD motifs are highlighted in green. A conserved sequence adjacent to the second HKD motif is highlighted in purple. The arrows indicate the point mutations introduced in this study.

To determine the contribution of individual amino acids in the HKD motifs of *T. brucei* TbCls to growth and CL synthesis, we first replaced histidine (H205N) or lysine (K207R) in the first HKD motif. Second, we generated TbCls enzymes containing point mutations in both HKD motifs (K207R+K542A) or in a conserved region adjacent to the second HKD motif (G494R). Conditional TbCls knockout parasites cultured in the presence of tetracycline to maintain expression of TbCls-HA grew logarithmically with a cell doubling time of 12 h (Fig. 2A), confirming previous findings(26). In contrast, in the absence of tetracycline, parasite growth slowed down after 24 h in culture, followed by a complete growth arrest after 48 h (Fig. 2A). This growth phenotype could be fully rescued by constitutive expression of wild-type TbCls-cMyc in the conditional TbCls knockout background (Fig. 2B). In contrast, expression of cMyc-tagged TbCls enzymes carrying point mutations in the first, second, or both HKD motifs were unable to restore normal growth of parasites (Figs 2C-F). Complete growth arrest was observed after replacing the histidine residue in the first HKD domain (H205N) (Fig. 2C), whereas replacement of the lysine residue in the same motif (K207R) only resulted in reduced growth (Fig. 2D). In the double mutant where the lysine residues in both HKD motifs were replaced (K207R+K542A) again complete growth inhibition was observed (Fig. 2E). Similarly, growth was completely arrested in parasites expressing TbCls-cMyc containing a point mutation (G494R) in a conserved region near the second HKD motif (Fig. 2F). In control experiments, re-expression of TbCls-HA in two selected cell lines (H205N and G494R) by addition of tetracycline to the culture medium at day 3 restored normal growth of parasites (Fig. 2G, H).

**Fig. 2.**
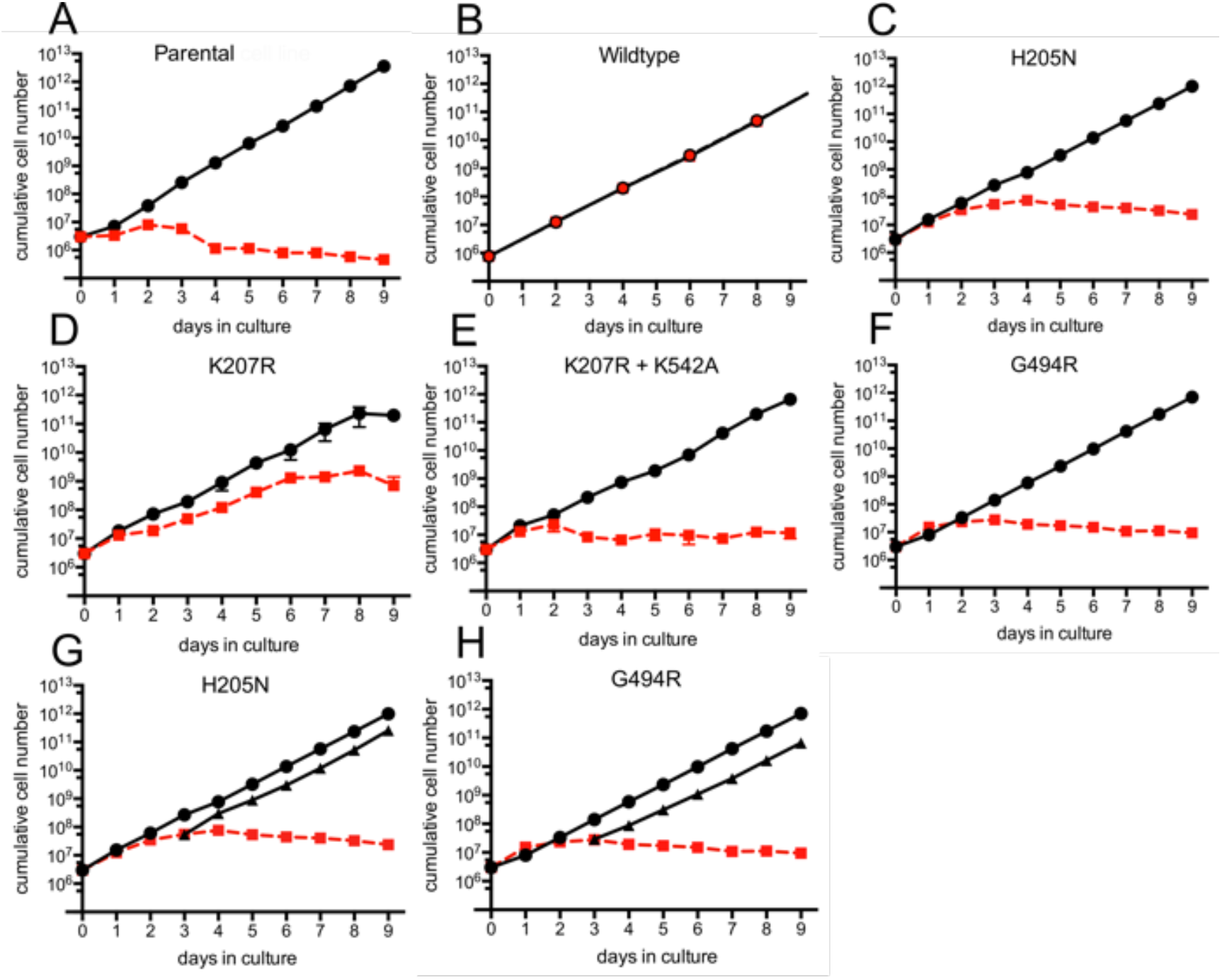
Growth of *T. brucei* procyclic forms. Parental parasites (A) and parasites constitutively expressing wild-type TbCls-cMyc (B) or mutant forms of TbCls-cMyc (C-H; mutations are indicated) were grown for 9 days in standard culture medium and the cell densities were recorded. The data points represent cell numbers (mean values ± standard deviations from 3 independent experiments) of parasites cultured in the presence (black circles) or the absence (red circles) of tetracycline to maintain or ablate, respectively, the expression of the inducible copy of TbCls-HA. In G and H, tetracycline was added to the cultures grown in the absence of tetracycline for 3 days to induce re-expression of TbCls-HA (black triangles). For some time points, the size of the symbol is larger than the standard deviation. In panel B, the data points for cells cultured in the presence and in the absence of tetracycline overlap.

In all parasite cell lines, expression of the mutated forms of TbCls-cMyc, as well as the tetracycline-dependent copy of TbCls-HA, was analyzed by immunoblotting using anti-cMyc and anti-HA antibodies, respectively (Fig. 3). The results show that TbCls-HA was expressed in all cell lines cultured in the presence of tetracycline and TbCls-cMyc was present in all cell lines except in parental parasites (Fig. 3A). In contrast, in parasites cultured in the absence of tetracycline, TbCls-HA was absent from all cell lines whereas TbCls-cMyc was expressed at similar levels in all parasites harboring constitutive wild-type or mutant TbCls-cMyc constructs (Fig. 3B).

**Fig. 3.**
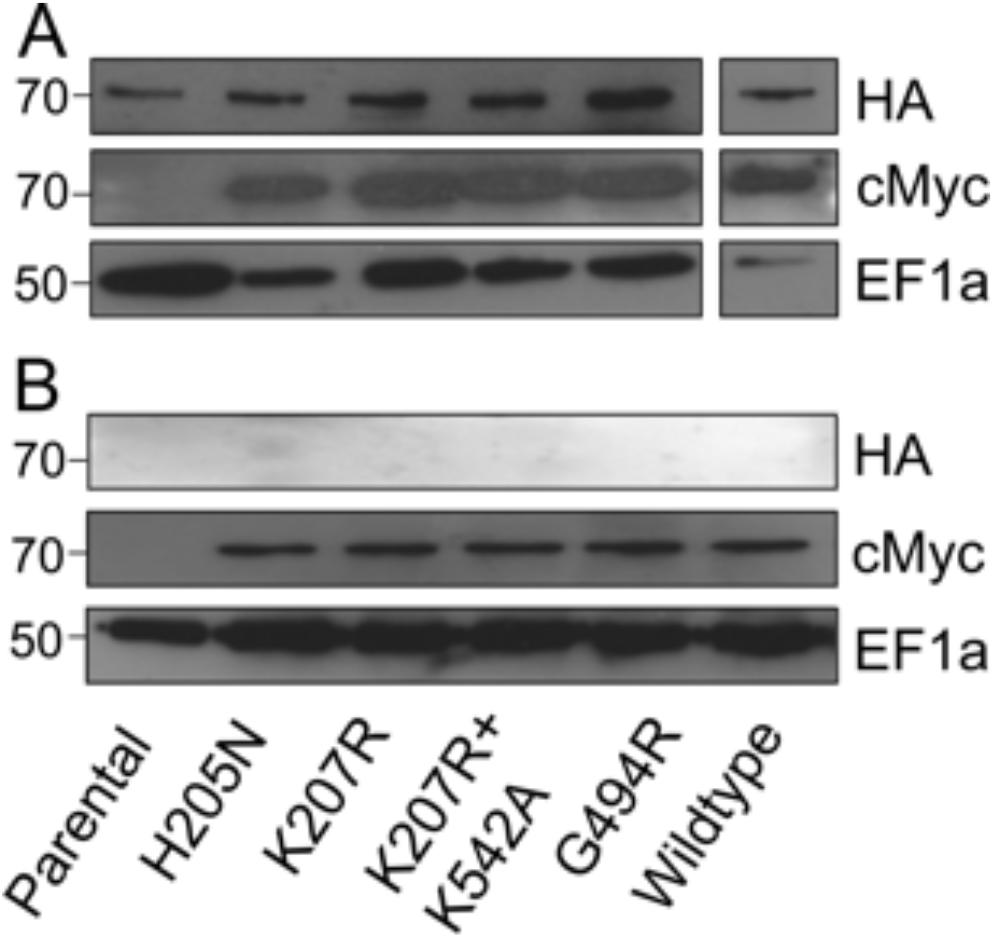
Immunoblot analysis of HA-tagged wild-type TbCls (TbCls-HA) and c-Myc-tagged mutant forms of TbCls (TbCls-cMyc). Whole cell lysates were prepared from trypanosomes cultured in the presence (A) or the absence (B) of tetracycline and separated by SDS-PAGE. After transfer to membranes, HA- and cMyc-tagged TbCls was detected using anti-HA and anti-cMyc antibodies, respectively, in combination with the corresponding fluorescent secondary antibodies. Eukaryotic elongation factor 1a (EF1a) served as loading control and was visualized using anti-EF1a primary antibody. TbCls-cMyc mutations are indicated in each lane; wildtype refers to unmutated cMyc-tagged TbCls. Molecular mass markers (in kDa) are indicated in the left margins.

Immunofluorescence microscopy revealed that all cMyc-tagged TbCls enzymes co-localized with *T. brucei* ATOM(28, 29), a marker protein of the outer mitochondrial membrane (Fig. 4). Together, these results demonstrate that all (mutant) TbCls-cMyc enzymes were constitutively expressed at similar levels in the conditional TbCls knockout background and localized properly to the mitochondrion.

**Fig. 4.**
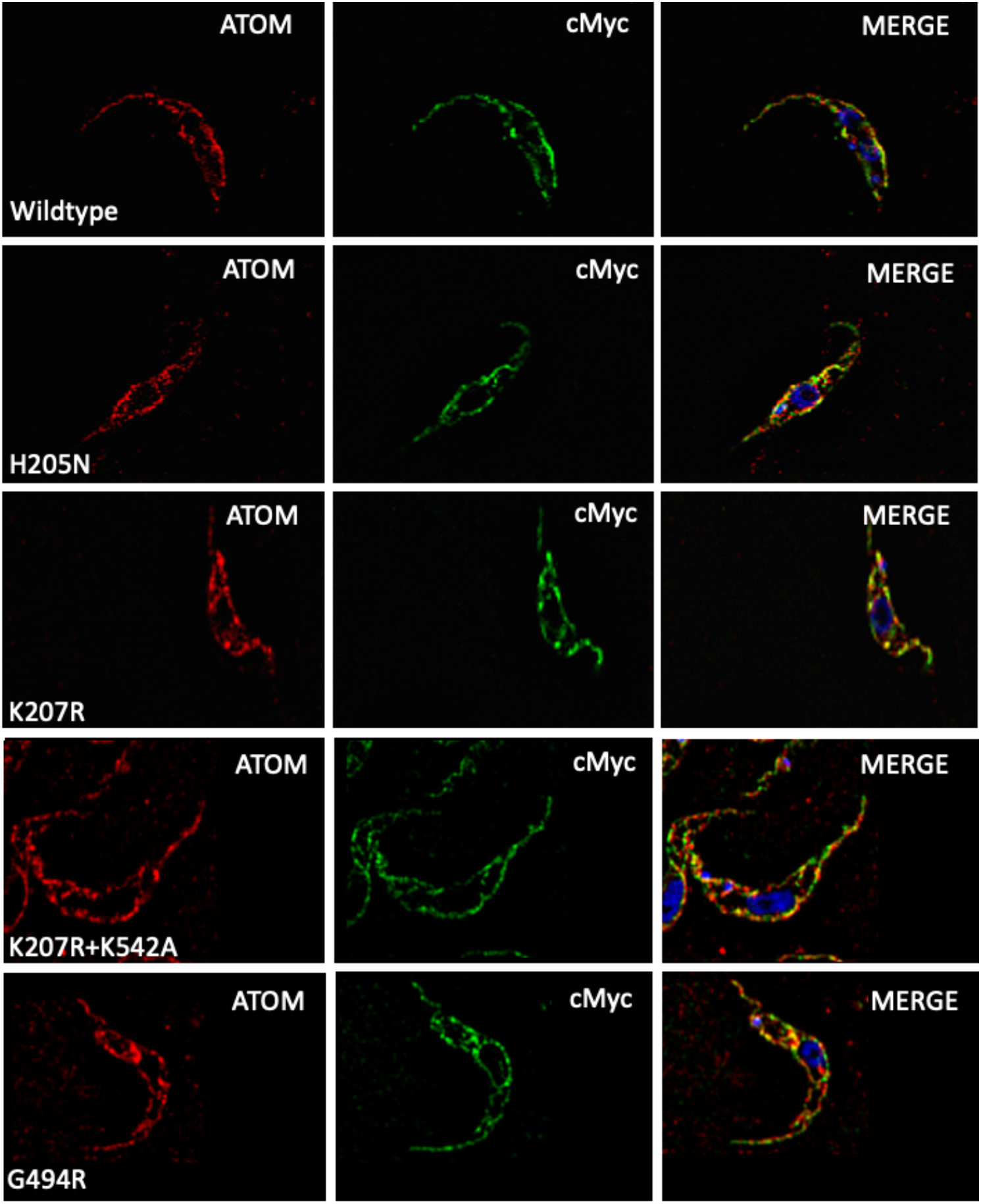
Localization of TbCls-cMyc enzymes by immunofluorescence microscopy. Parasites expressing c-Myc-tagged wildtype or mutant forms of TbCls were co-stained with antibodies against c-Myc (middle panels) and the mitochondrial marker ATOM (left panels). The right panels show the merge images. DNA was visualized by DAPI in blue (scale bar = 5 μm).

### Point mutations in the HKD motif reduce *de novo* CL synthesis

To study *de novo* synthesis of CL in *T. brucei* procyclic forms, parasites expressing wild-type or mutated forms of TbCls-cMyc were cultured in the presence or absence (to ablate TbCls-HA expression) of tetracycline for three days and phospholipids were labeled *in vivo* by adding [^3^H]-glycerol to the trypanosomes in cultures. This results in incorporation of radioactivity into all glycerophospholipid classes in *T. brucei* procyclic forms, including CL(26, 27). After incubation, labeled lipids were extracted, separated by thin-layer chromatography (TLC) and analyzed by radioisotope scanning. In line with a previous report(26), we show that labeling of parental parasites cultured in the presence of tetracycline, to maintain expression of TbCls-HA, resulted in the synthesis of [^3^H]-labeled PG and CL (Fig. 5A). In contrast, after ablation of TbCls-HA, [^3^H]-CL production was completely inhibited (Fig. 5A). In addition, we show that the formation of [^3^H]-CL was restored completely in conditional TbCls knock-out parasites expressing wild-type TbCls-cMyc cultured in the absence of tetracycline (Fig. 5B), demonstrating that the constitutively expressed wild-type enzyme is fully active. Restoration of [^3^H]-CL formation by wild-type TbCls-cMyc was not dependent on the labeling time with [^3^H]-glycerol, however, the relative amounts of radioactivity incorporated into CL and PG differed considerably (Fig. 6A, B). After 4 h of labeling with [^3^H]-glycerol, approximately 6% and 12% of total radioactivity in the glycerophospholipid fraction was incorporated into CL and PG, respectively (Fig. 6A), whereas after 16 h of labeling the relative amount of radioactivity in CL increased by more than 4-fold (to approximately 27%), while that in PG remained largely unchanged (at approximately 10%) (Fig. 6B). The relative increase in [^3^H]-CL formation after extended labeling time is consistent with PG being a precursor for CL synthesis and the reported slow turnover of CL(30, 31).

**Fig. 5.**
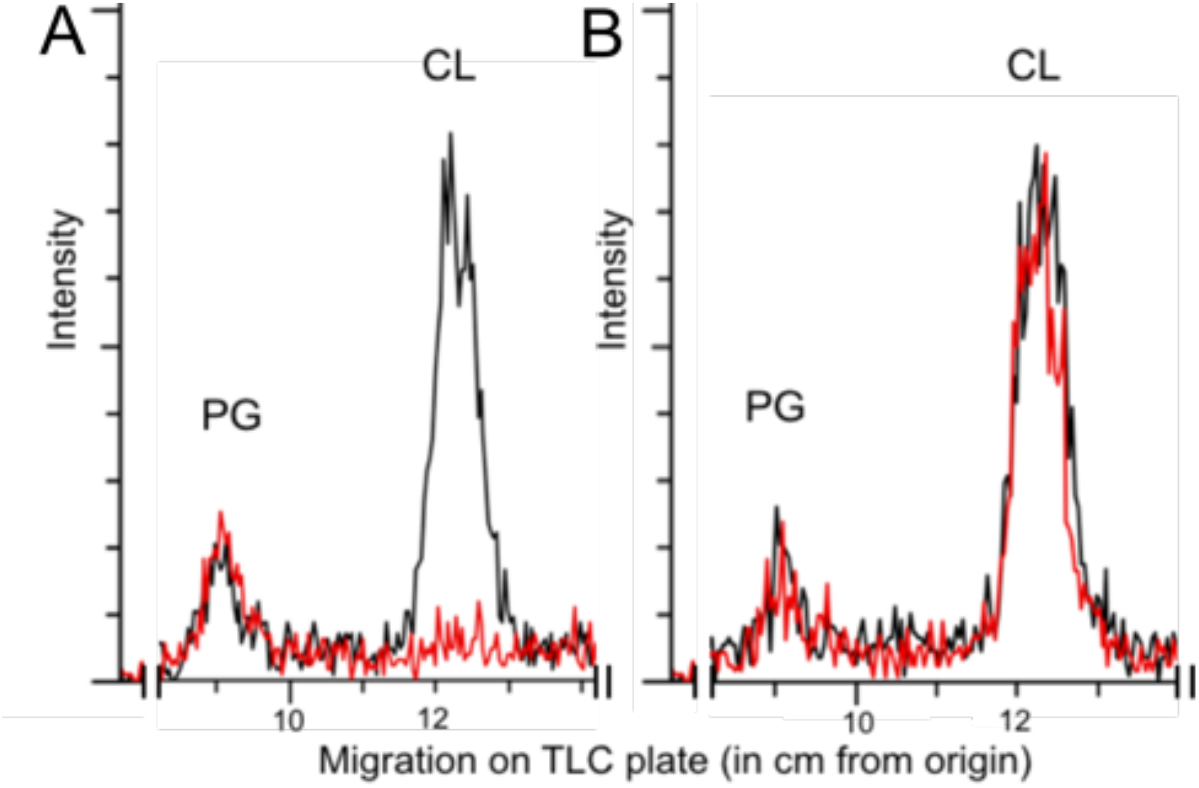
Analysis of *de novo* PG and CL synthesis. Parasites were cultured in the presence (black) or the absence (red) of tetracycline to maintain or ablate, respectively, TbCls-HA expression and incubated with [^3^H]-glycerol for 16 h. Phospholipids were extracted and separated by TLC and incorporation of radioactivity into the different phospholipid classes was analyzed by radioisotope scanning. Only the traces of *de novo* synthesized [^3^H]-PG and [^3^H]-CL in parental cells (A) and parasites expressing wildtype TbCls-cMyc (B) are shown.

**Fig. 6.**
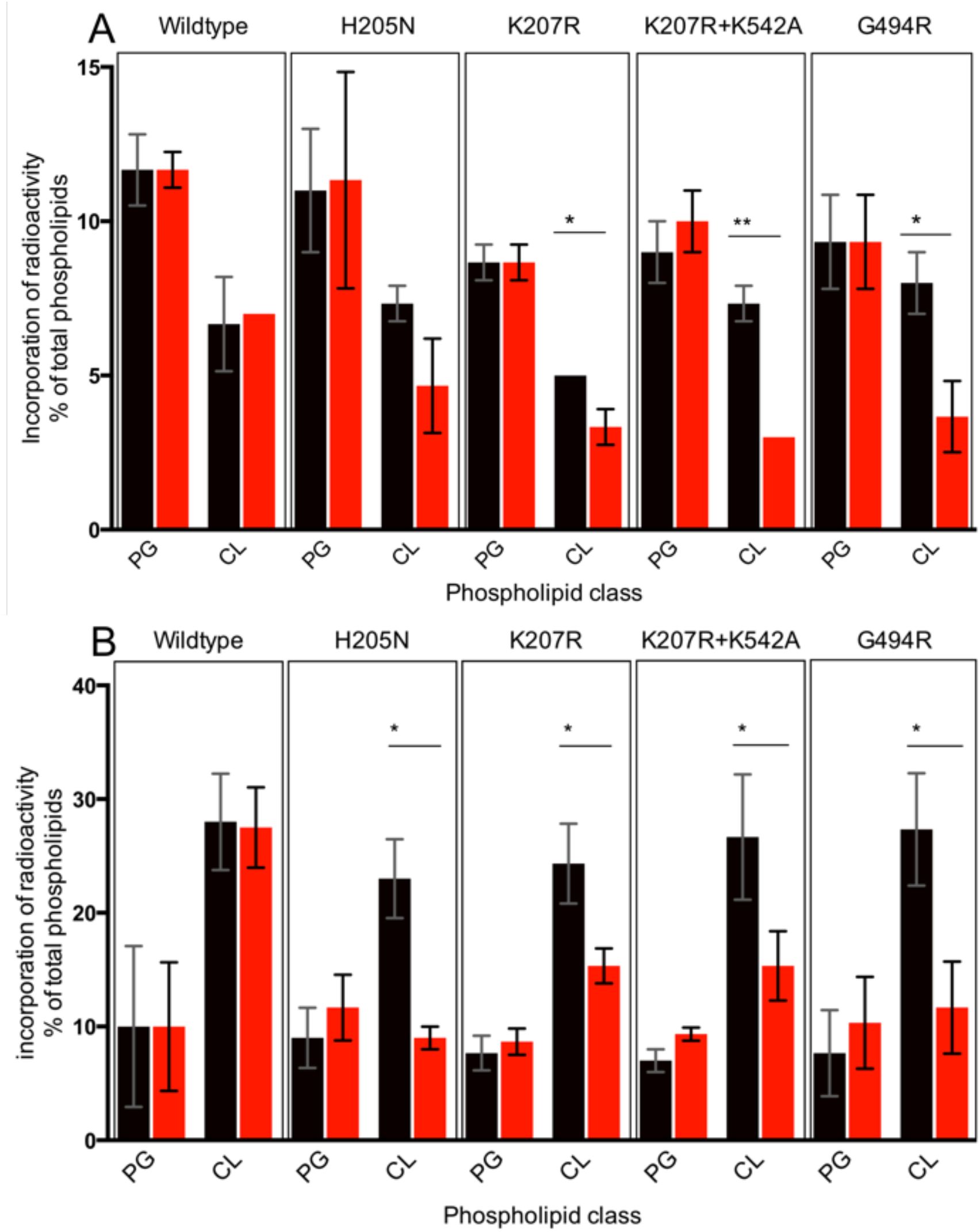
[^3^H]-PG and [^3^H]-CL formation in parasites expressing mutant TbCls-cMyc. Trypanosomes were cultured in the presence (black) or the absence (red) of tetracycline to maintain or ablate, respectively, TbCls-HA expression and incubated with [^3^H]-glycerol for 4 h (A) or 16 h (B). Phospholipids were extracted and separated by TLC and incorporation of radioactivity into PG and CL was quantified by radioisotope scanning. Data are from at least 3 independent experiments (means ± standard deviations). The asterisks indicate statistically different values (determined using t-tests) between parasites cultured in the presence or the absence of tetracycline.

In parasites expressing mutant forms of TbCls-cMyc, production of [^3^H]-CL was inhibited (Fig. 6A, B). Incorporation of radioactivity into CL was reduced by 20-60% in all parasites expressing mutated TbCls-cMyc enzymes after 4 h of labeling with [^3^H]-glycerol (Fig. 6A). No changes were observed in the formation of [^3^H]-PG (Fig. 6A). Similar results were obtained after 16 h of [^3^H]-glycerol labeling, with reductions in [^3^H]-CL formation of 40-60% compared to cells expressing TbCls-HA (Fig. 6B). Although incorporation of label into PG after 16 h of labeling was consistently higher in extracts obtained from parasites cultured in the absence of tetracycline compared to cells cultured in the presence (Fig. 6B), the differences were statistically not significant.

In summary, we used *T. brucei* procyclic form parasites, in which the expression of the essential wild-type TbCls can be ablated, to study CL synthesis by TbCls enzymes carrying point mutations in the two conserved HKD motifs characteristic for PLD-type enzymes. Conservative substitutions in either of the HKD motifs resulted in inhibition of *de novo* synthesis of CL and reduced growth of parasites, while non-conservative mutations resulted in inhibition of CL formation and parasite proliferation. This study highlights the importance of both HKD motifs, and selected residues in a conserved region flanking the second HKD motif, of TbCls in the biosynthesis of CL and parasite survival in culture.

## Acknowledgements

We are grateful to Petra Gottier for providing help with the TbCls add-back plasmids. The work was supported by grant 169355 from the Swiss National Science Foundation to P.B.

## Materials and Methods

### Reagents

All reagents were of analytical grade and purchased from Merck-Sigma Aldrich (Darmstadt, Germany) or MilliporeSigma (Burlington, MA, USA) unless otherwise stated. Kits for plasmid DNA extractions, PCR purifications, vector dephosphorylation and ligation reactions were purchased from Promega (Wallisellen, Switzerland). All sequencing was done by Microsynth AG (Balgach, Switzerland).

### Trypanosome cultures

Trypanosomes were cultured at 27 °C in SDM79 (Invitrogen, Basel, Switzerland) supplemented with 10% heat inactived fetal bovine serum (Thermo Fisher Scientific, Reinach, Switzerland) and a mixture of hemin and folic acid (160 µM hemin + 90 µM folic acid). *T. brucei* TbCls conditional knock-out (29-13) procyclic forms were cultured in the presence of hygromycin (25 μg/ ml), G418 (15 μg/ ml), puromycin (2 μg/ ml), blasticidin (5 μg/ ml), phleomycin (0.2 μg/ ml) and tetracycline (1 μg/ ml). Growth media for parasites constitutively expressing mutated or wildtype TbCls in addition contained nourseothricin (1.5 μg/ ml).

### Mutagenesis

QuickChange site-directed mutagenesis kit (Agilent Technologies, Basel, Switzerland) was used to carry out single and double point mutations in expression plasmids. Primers used to substitute targeted residues are listed in the supplemental data. Plasmids were sequenced to confirm the mutations and the unchanged sequences of the adjacent regions.

### Whole cell lysates and crude membrane preparations

For whole cell lysate preparations, 10^7^ trypanosomes were harvested by centrifugation and the resultant pellet was washed with Tris-buffered saline (TBS, 10 mM Tris-HCl, pH 7.5, 144 mM NaCl), resuspended in sample buffer (2.5% SDS) and incubated at 50 °C for 5 min.

For crude membrane preparations, a digitonin extraction was performed. Trypanosomes (10^8^ cells) were harvested by centrifugation and washed in TBS before being resuspended in 0.5 ml buffer (20 mM Tris-HCl, pH 7.5, 600 mM sorbitol, 2 mM EDTA). This was followed by addition of the same buffer supplemented with 0.05% (w/v) digitonin. After 5 min incubation on ice, the suspension was centrifuged at 6000 x g. Subsequently, 100 μl extraction buffer (20 mM Tris-HCl, pH 7.2, 15 mM NaH_2_PO_4_, 0.6 M sorbitol) containing 1.5% (w/v) digitonin was added to the pellet. The sample was incubated on ice for 15 min and membranes were pelleted by centrifugation at 16’000 x g in a tabletop centrifuge. The resultant pellet was used for further analysis.

### SDS PAGE

Proteins were denatured by SDS and run on a 10% polyacrylamide gel as described previously(32). The proteins were then transferred onto nitrocellulose membranes (Thermo Fisher Scientific) using a semidry protein blotting system (Bio-Rad, Hercules, CA, USA) for 75 min at a maximal current of 2.5 mA cm^−2^ gel area. The membranes were blocked for 1 h in TBS, containing 5% (w/v) milk powder, and proteins of interest were identified using specific primary and horseradish peroxidase-conjugated secondary antibodies. Protein-specific antibodies were visualized using SuperSignal™ West Pico PLUS Chemiluminescent Substrate (Thermo Fisher Scientific). The following antibodies were diluted in TBS, containing 5% milk: mouse monoclonal anti-cMyc (9E10; Santa Cruz Biotechnology, Dallas, TX, USA; dilution 1:1000); mouse monoclonal anti-HA (HA11; Enzo Life Sciences, Farmingdale NY, USA; dilution 1:3000); mouse anti-EF1a (Covance, Fullinsdorf, Switzerland; dilution 1:20000); HRP-conjugated rat anti-mouse antibody (DAKO-Agilent, Basel, Switzerland; dilution 1:5000); HRP-conjugated goat anti-rat antibody (Thermo Fisher Scientific, dilution 1:10000); rabbit anti-Cox4 and mouse anti-Hsp70 antibodies (kindly provided by A. Schneider, University of Bern, Bern, Switzerland; dilution 1:1000).

### Immunofluorescence microscopy

Trypanosomes (10^6^ cells) were harvested by centrifugation, resuspended in ice cold PBS (phosphate-buffered saline; 137 mM NaCl, 2.7 mM KCl, 10 mM Na_2_HPO_4_, 2 mM KH_2_PO_4_, pH 7.4), allowed to adhere onto glass slides (Thermo Fisher Scientific) and fixed with 4% (w/v) paraformaldehyde for 10 min. Fixed cells were washed with PBS, permeabilized with 0.2% (w/v) Triton X-100 for 5 min followed by blocking in PBS, containing 2% (w/v) bovine serum albumin, for 30 min, and incubated with antibodies diluted in blocking solution for 45 min. The following antibodies were used: mouse monoclonal anti-cMyc (9E10; Santa Cruz Biotechnology; dilution 1:250); rabbit anti-ATOM (kindly provided by A. Schneider, University of Bern, Bern, Switzerland; dilution 1:1000). After 3 washes in PBS for 5 min each, the slides were incubated with the corresponding secondary fluorophore-conjugated antibodies: Alexa Fluor goat anti-mouse 488 and goat anti-rabbit 594 (Thermo Fisher Scientific; dilution 1:1000) in blocking solution for 45 min. Finally, samples were washed, air dried and mounted with Vectashield (Vector Laboratories, Burlingame, CA, USA) containing 4’,6-diamidino-2-phenylindole (DAPI).

Fluorescence microscopy was performed with a Leica DM 16000 B inverted microscope using a 60x oil objective and images were acquired using Leica DFC360 FX camera. Image deconvolution and processing was performed using Leica LAS X and Fiji software (National Institutes of Health).

### [^3^H]-glycerol labeling and lipid analysis

Trypanosomes in culture (10^8^ cells) were labeled with 10 μCi [^3^H]-glycerol (Anawa, Kloten, Switzerland) for 4 h or 16 h before being harvested by centrifugation and washed twice with TBS. The radiolabeled lipids were extracted according to Bligh and Dyer(33), spotted onto silica gel plates (Merck, Zug, Switzerland) and separated by one dimensional TLC using a solvent system consisting of chloroform: methanol: acetic acid (65: 25: 8, by vol). The plates were dried, and radioactivity was detected using a radioisotope scanner (Berthold Technologies, Bad Wildbad, Germany). Data processing was done using the Rita Control software provided by the manufacturer.

## Supplemental data: List of primers used to mutagenize TbCls

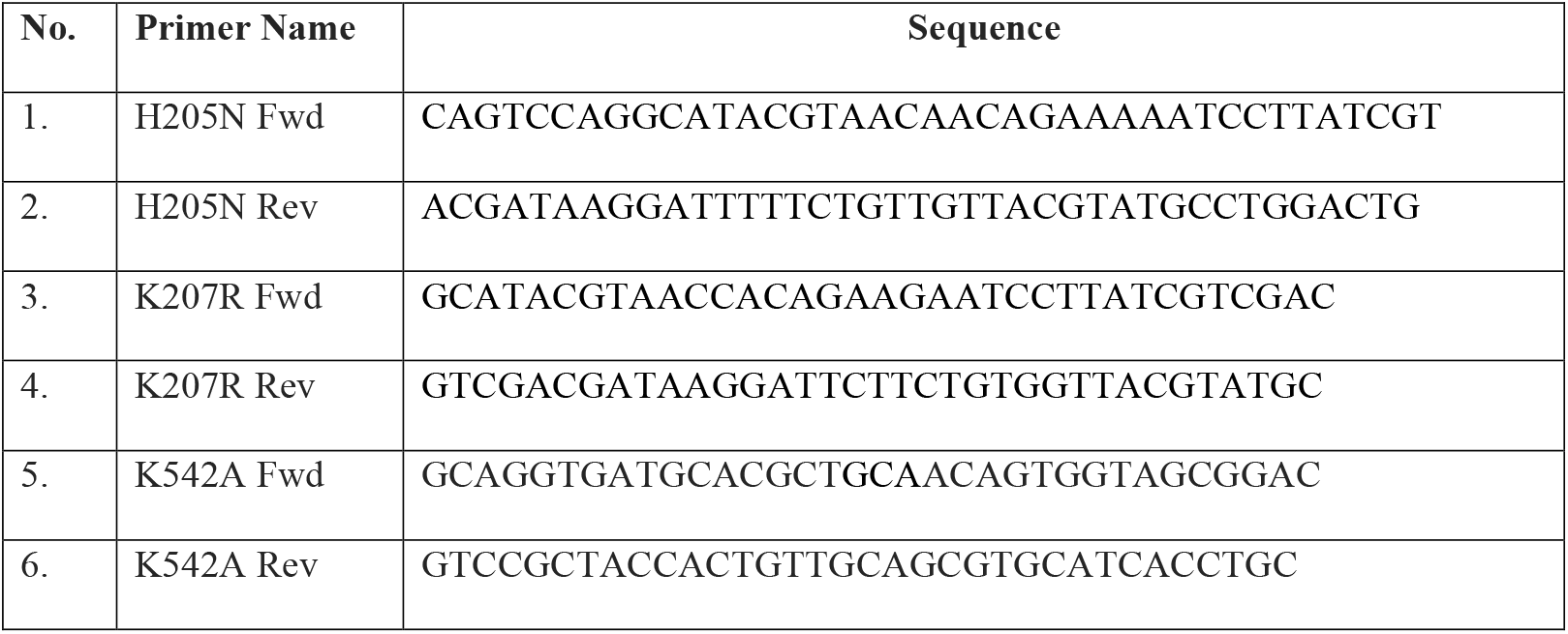
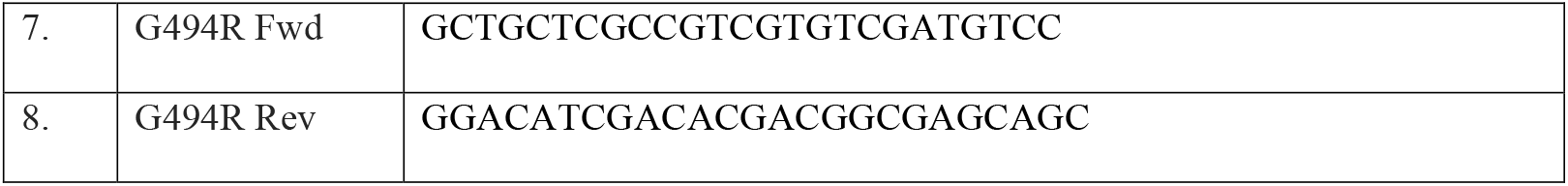

